# Beyond compliance: evaluating the impact of assent-based training on equine behavior and the human-horse relationship

**DOI:** 10.64898/2026.09.12.751177

**Authors:** Elisa Tanghetti, Remco Folkerstma, Friederike Range

## Abstract

Recent advances in animal training increasingly emphasize welfare, positive experience and the reduction of stress. In the equine sector, however, such progress is complicated by the horse’s dual role as both companion and performance animal. This study investigated whether an assent-based training approach (ABT) has an effect on horse behavior and human–horse interaction quality compared to traditional equine training (TET) based on pressure and release. Two groups of horse–owner pairs (ABT and TET) participated in two behavioral tests. A stroking session evaluated horses’ responses to human tactile interaction, while an obstacle path assessed horses’ exploration of novel objects and refusal behaviors. Behavioral measures of exploration, avoidance, and owner-directed behaviors were analyzed alongside horse personality traits. A newly validated questionnaire was used to gather demography information, training habits and horse personality traits. During the stroking session, ABT horses displayed more positive human-directed behaviors, reflecting active engagement rather than passive tolerance. Sociability was associated with the overall frequency of human-directed behaviors, but did not predict their positive valence, which was instead linked to ABT training style. Horses trained mainly with ABT explored obstacles more and showed fewer refusal behaviors than traditionally trained horses, independently of anxiety traits derived from personality scores. These findings suggest that assent-based training practices are associated with increased confidence and cooperative engagement in horses, supporting the potential of agency-oriented approaches to enhance both equine welfare and human–horse interaction quality.

## Introduction

In recent decades, human–animal relationships (HARs) have evolved alongside advances in animal welfare science and a growing recognition that animals experience not only suffering but also positive states of well-being. This shift has encouraged training and management practices aimed at reducing fear, stress and coercion while promoting voluntary engagement. Within this broader context, horses represent a particularly complex case due to their liminal status as both companion and performance animals, exposing them to unique welfare challenges not faced by other domestic species (Carroll, 2022).

Traditional equine training (TET) largely relies on negative reinforcement through pressure-and-release techniques, often supplemented by punishment to suppress unwanted behaviors (DeAraugo, 2014). While these methods can produce a reliable task completion, their effectiveness depends on precise timing. Inconsistent or delayed release of pressure may lead to confusion, fear and conflict behaviors such as head tossing, rearing or avoidance (Goodwin, 2009). Horses’ resistance in such cases is usually framed as disobedience, reinforcing cultural narratives that justify punishment and pushing horses beyond their physical or cognitive limits (Goodwin, 2009; McLean, 2017). These practices raise concerns regarding both learning efficiency and equine welfare.

In response, alternative approaches such as natural horsemanship have emerged, promoting the idea that humans can “speak horse” by mimicking equine social signals to establish leadership (DeAraugo, 2014). Although these methods emphasize attentiveness to the horse’s perspective, they remain primarily grounded in negative reinforcement and avoidance of pressure. Moreover, dominance-based interpretations of horse behavior lack strong ethological support. Morphological and perceptual differences make it unlikely that horses perceive humans as conspecifics, undermining assumptions that interspecific interactions replicate herd dynamics (Henshall, 2014). Empirical evidence further indicates that dominance rank in horses is not a fixed trait but a context-dependent relationship, and that leadership within groups is situational and distributed rather than tied to dominance or resource control (Drews, 1993; Hartmann, 2017; Van Vugt, 2008; Bourjade, 2015). Thus, the horses trained with natural horsemanship likely learn primarily through the release from pressure rather than recognition of human social rank (Hartmann, 2017).

More recently, training approaches grounded in learning theory have gained attention, emphasizing systematic use of positive reinforcement. Positive reinforcement, widely applied in companion and zoo animal training, involves rewarding desired behaviors immediately after they occur and has been associated with increased motivation, problem-solving behavior, and voluntary engagement (Innes & McBrided, 2008). In horses, using positive reinforcement has been linked to increased contact-seeking behavior, though findings regarding emotional state and stress physiology remain mixed (Larssen, 2022). Concerns include heightened arousal, frustration-related behaviors, and potential aggression if reinforcement is inconsistently applied (McLean, 2017; Kieson, 2020). Nonetheless, when implemented with appropriate timing and criteria, positive reinforcement may reduce reliance on aversive stimuli and support improved welfare outcomes.

Building on learning theory, voluntary participation protocols have been increasingly adopted in companion and zoo animal contexts, where animals are allowed to actively opt in or out of procedures (Dadone, 2016). These protocols acknowledge animals’ capacity to learn choice contingencies and respect their ability to refuse participation. Evidence suggests that horses can use symbolic cues to communicate preferences in free-choice situations (Medjell et al., 2016). Assent-based training (ABT) extends this principle by teaching a specific signal, typically through positive reinforcement, allowing horses to indicate readiness to participate and to withdraw assent by disengaging (Innes, 2008; Carroll, 2022; Wess, 2022). Such approaches are proposed to increase predictability and perceived control, reducing defensive behaviors and improving both horse welfare and human safety.

Despite growing interest in assent-based training, empirical evidence evaluating their effects on the human - horse relationship (HHR) relative to traditional equine training remains scarce. It is unclear whether granting horses greater control influences their willingness to engage in novel or emotionally salient contexts, or whether such methods can achieve outcomes comparable to conventional approaches. Importantly, evaluating training effects must account for individual differences that can shape behavioral responses during human–horse interactions and therefore require systematic consideration. Horse personality traits, such as sociability and anxiousness, have been shown to affect responses to novelty, handling and human contact, shaping reactivity and communicative expression (Lloyd et al., 2008; Carroll et al., 2022). Finally, owner behavior may influence the animal’s behavior via the owner’s emotional state (Keeling 2009; Samhita 2013), attachment style or level of education (DeAraugo, 2014).

The present study tested the hypothesis that horses trained mainly using assent-based training (ABT) differ from horses trained using traditional equine techniques (TET) in regard to human directed behaviors and exploration. We used two behavioral tests – a stroking session and an obstacle path - to assess horses’ behaviors, while controlling for horses’ sociability and anxiousness. For the **stroking session**, we predicted that horses trained with ABT would display more positive human-directed behaviors, reflecting greater engagement during tactile interaction, while the overall frequency of aversive behaviors would not differ markedly between groups. We further predicted that sociability would be associated with a higher frequency of positive human-directed behaviors. For the **obstacle path**, we predicted that horses trained with ABT would show greater explorator y behavior and fewer refusal behaviors compared to TET horses. We further predicted that anxiousness would be associated with reduced exploration and increased refusal behaviors. Finally, we investigated the human behavior in the challenging situation to account for potential differences of owners’ interactions expecting that owners practicing ABT methods would use more praise and less pressure than owners practicing TET in accordance with the training philosophies.

## Methods

### Ethics

All experimental protocols were approved by the relevant institutional ethics and animal welfare committees. The pilot studies conducted in Austria were approved by the Ethics and Animal Welfare Committee of the University of Veterinary Medicine Vienna (ETK protocol No. 128/11/2024). The behavioral tests conducted in Italy were approved by the Animal Welfare Committee of the University of Bologna (ID 4095; Protocol No. 172870/2025). The survey component received a favorable ethical evaluation from the Ethics Committee of FH Campus Wien, which issued a waiver on 14 May 2025.

All methods were carried out in accordance with the relevant institutional and national guidelines and regulations applicable in Italy and Austria. Written informed consent was obtained from all owners for their participation with their animals in the study and for the publication of the images and data included in this article.

### Subjects

A total of 45 horse–owner pairs participated in the behavioral study. Horses were assigned to one of two groups based on preliminary phone interviews. The assignment was later confirmed based on information reported by the owners on their main activity and training philosophy in the survey (see below). The ABT group consisted of 24 horses (13 females, 11 geldings; age average: 13; age range: 7-25) trained using methods characterized by assent-based protocols, voluntary participation and systematic use of positive reinforcement. The ABT group participants had participated in courses inspired by Co-Creational Horsemanship (Ghizzoni, G. Home, https://hathaequus.com/, accessed on August 2024), an emerging equine training approach developed at the Wildsong Ranch, Colorado, that integrates learning theory, ethical considerations and principles from ecological dynamics (Brando, 2023; Davies, 2023). Co-Creational Horsemanship minimizes the use of negative reinforcement outside of safety requirements and emphasizes positive reinforcement alongside structured assent protocols.

The comparison group (TET) included 21 horses (10 females, 10 geldings, 1 stallion; age average: 12, age range: 5 - 30) trained using conventional or conspecific-based approaches relying primarily on pressure-and-release and punishment. Both groups consisted of mixed and pure breeds to the same extent (see supplementary Table S1).

To minimize variability due to confounding factors, inclusion criteria ensured comparable living and management conditions across groups: most horses (39/45) spent most of the day in paddocks and were socially housed, 5 TET horses were given access to paddocks only in good weather conditions and 1 TET horse had never access to paddocks. All owners were female. Four owners participated with two horses and one with three horses, all the other owners participated with just one horse. We included only participants that had owned/taken care of their horse (42 owners, 3 main care takers) for at least one year and worked with them at least twice per week. An a priori power analysis indicated that a minimum of 15 horse–owner pairs per group was sufficient to detect medium-sized effects with 80% power at a significance level of 0.05; a larger sample was tested to account for potential dropouts.

### Questionnaire

To assess demographic variables, horse personality traits and owner’s training practices, an online questionnaire was developed based on published studies that the owners had to complete within a month after participating in the study.

Items investigating horse sociability and anxiousness were adapted from the Horse Personality Questionnaire (HPQ) of Lloyd et al. (2008). For each item, respondents indicated their likelihood on a five-point Likert scale ranging from very unlikely to very likely. Responseswere coded using a symmetric scoring scheme from −2 (very unlikely) to +2 (very likely) (see Supplementary Table S2). Personality trait scores were calculated by summing the scores of the items associated with each trait (sociability and anxiousness), with higher scores indicating a stronger expression of the respective trait.

As mentioned above, participants were initially assigned to ABT or TET groups based on a preliminary phone interview assessing their training practices. To validate that this classification accurately reflected owners’ training approaches, a questionnaire section investigating different training concepts was included. Items assessed the use of positive reinforcement (R+), negative reinforcement (R−), assent-based training (ABT), and traditional equine training techniques (TET), with R+ and R− items adapted from Lundberg et al. (2020) (see Supplementar y Table S3). Responses were collected on five-point Likert scales and converted into item-level scores using a symmetric coding scheme (0 to ±4), with R+ and ABT items scored positively and R− and TET items scored negatively. A composite training score (CTS) was calculated for each participant by summing all item-level scores, with higher values indicating greater alignment with R+/ABT practices and lower values indicating greater alignment with R−/TET practices. Mean CTS values were calculated for each group (ABT: mean = 21.5, SD = 8.03; TET: mean = −5.00, SD = 6.30), and the midpoint between group means (cut-off = 8.25) was used to assess consistency between questionnaire-based scores and initial classification. Two horse–owner dyads showed discrepancies between interview-based and questionnaire-based classification and were excluded from further analyses. Three participants did not complete the questionnaire; however, given the high consistency between interview and questionnaire data, these dyads were retained and classified based on the preliminary interview.

### Behavioral tests

Two behavioral tests were conducted to assess horses’ responses to human interaction and a challenging situation. The stroking session was designed to evaluate horses’ affective responses and engagement during prolonged human tactile interaction, whereas the obstacle path assessed horses’ willingness to explore novel stimuli, cross unfamiliar obstacles and their behavioral responses when guided by their owner in a potentially challenging situation.

#### Experimental setting

All behavioral tests were conducted outdoors at the horses’ home facilities, in familiar environments chosen by the owners to minimize contextual stress. Testing sessions took place in the afternoon under comparable weather conditions. Each horse–owner pair completed both tests within a single session lasting approximately 20 minutes. The stroking session was always conducted first, followed by the obstacle path. All sessions were video-recorded using a movable camera operated by an experimenter, ensuring full visibility of the horse’s body and audio recording for later behavioral coding.

#### Procedure

##### Stroking session

The stroking session lasted 8 minutes and took place in an area where the horse was accustomed to being groomed. Horses were loosely tied using halter and rope to allow limited freedom of movement. Owners were instructed to stroke the horse’s neck, head, back, belly, and hindquarters, adjusting pressure according to the horse’s responses. If the horse showed approach behavior toward a specific area, owners focused on that area with increased pressure; if the horse showed avoidance or no response, pressure was reduced and the area changed. After five minutes, owners continued the interaction using a brush following the same procedure for 3 minutes.

##### Obstacle path

The obstacle path was set up in a familiar arena or open space and consisted of three unfamiliar obstacles arranged sequentially (Figure 1): a 1.20 × 2 meters colorful blanket (Figure 2.a), a 1.5 meter wide passage with hanging balloons on the sides (Figure 2.b) and a 1.8 meter high archway with suspended plastic strips (Figure 2.c). For each obstacle, two poles placed three meters before and after the object marked the start and end of the trial (Figure 1). Horses were led on a loose rope by their owners and invited to cross each obstacle. Owners were instructed not to use treats, whips or any other tool. To cross each obstacle a maximum duration of 5 minutes was set to minimize prolonged discomfort; if the horse failed to cross within this time, the pair proceeded to the next obstacle. The use of three different obstacles provided repeated measures for each subject, reducing the influence of single-event responses potentially linked to past experiences.

**Figure 1:**
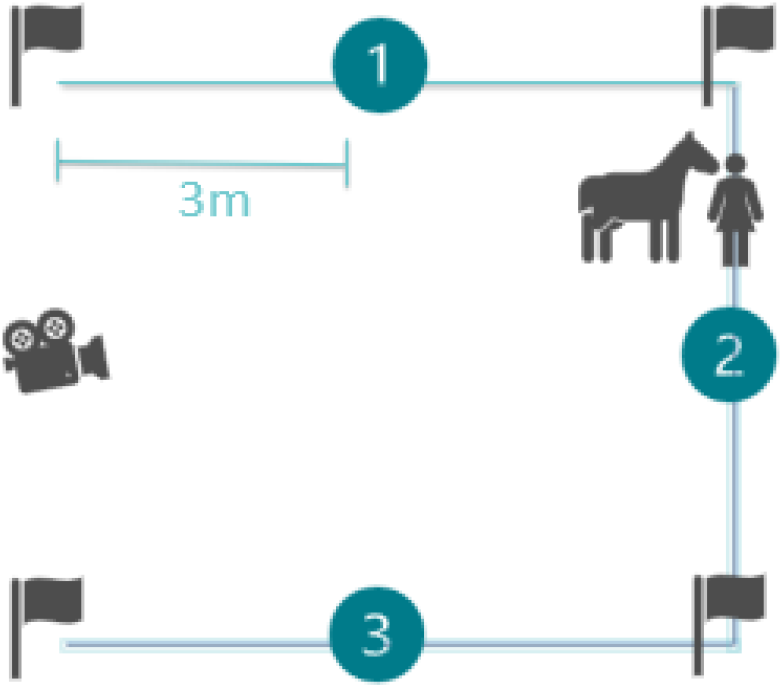
graphical representation of the Obstacle path experimental setting

**Figure 2:**
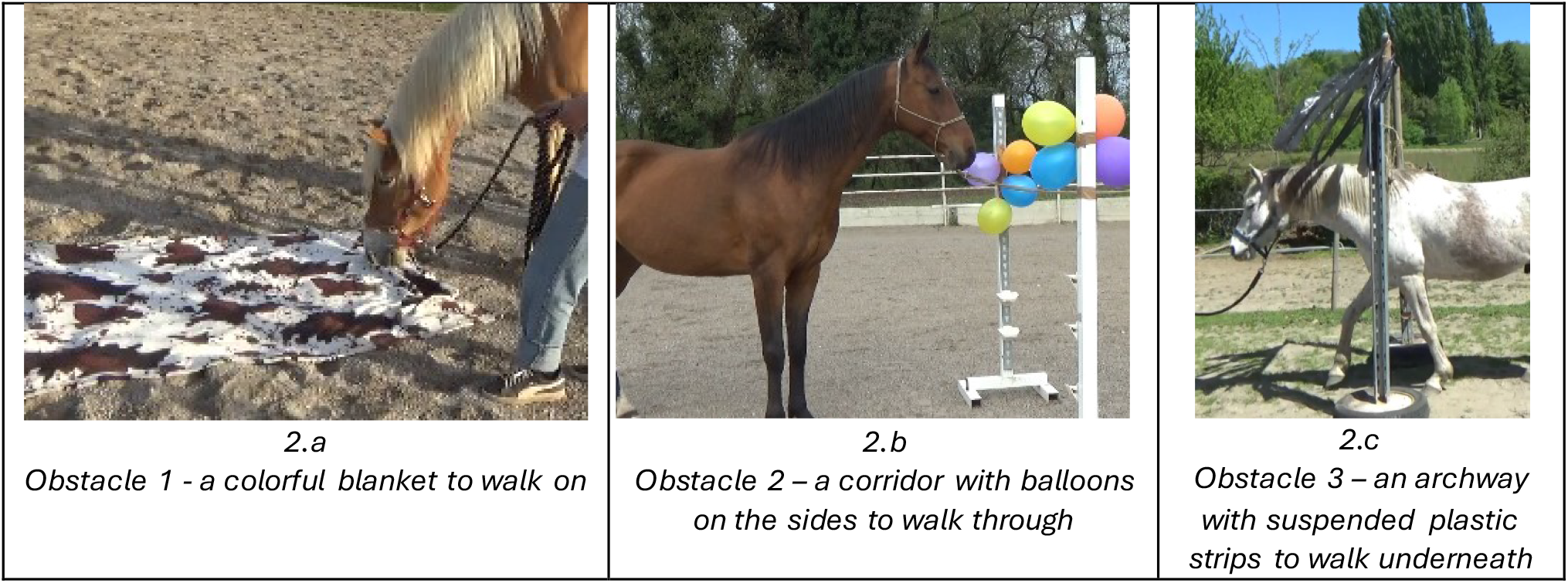
the 3 obstacles used in the Obstacle path test

**Figure 3:**
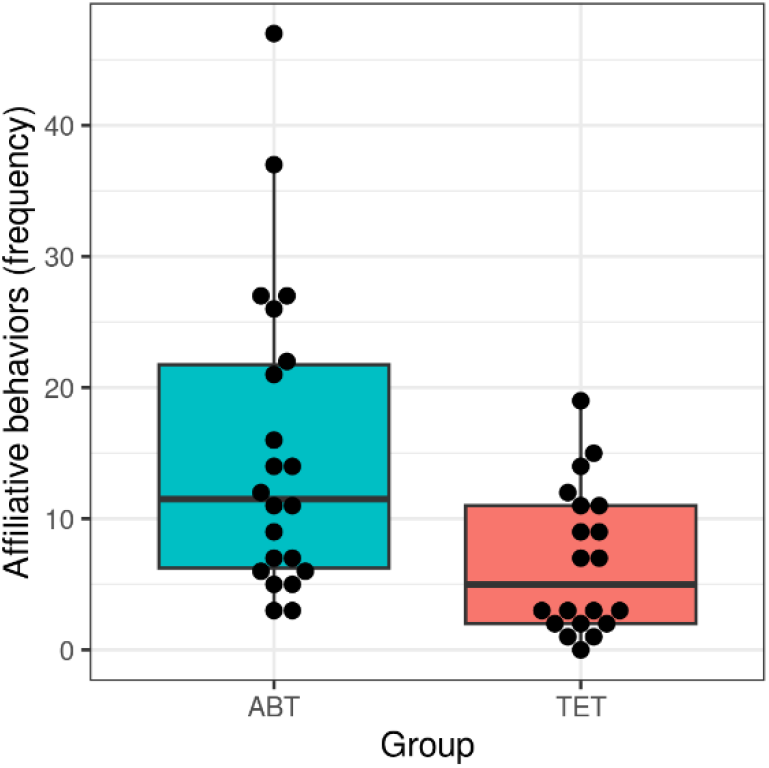
Frequency of affiliative behaviors during the stroking session in horses trained with assent-based (ABT) and traditional (TET) training protocols.

### Behavioral coding

Videos were coded using BORIS software v.8.25.4 (Friard & Gamba, 2016) according to predefined ethograms for each test. For the stroking session, we coded the duration of activity states (sleep, rest standing, alert, restless)(McDonnell, 2003), exploration of the brush, avoidance (threat, avoid, step aside and stomp) and affiliative (groom, scratch, look at and smell) behaviors. For the obstacle path, we coded the duration of crossing time, explorative behaviors, refusal behaviors (stationary and proactive)(Ijichi, 2008) and occurrence of owner behaviors (pulling on the rope, vocal praising, physical praising). To investigate the quality of crossing behavior, we measured the frequency of relaxed pass through, tense pass through, rush through and no pass (see Supplementary Tables S4 and S5 for definitions of behaviors). Inter-observer reliability was assessed on 20% of the tests (n = 10), independently coded by a second observer. Continuous behavioral measures were assessed by calculating the intraclass correlation coefficient using the function “icc” in the package “irr” (version 0.84.1) of the statistical program R (version 4.5.2) (R Core Team, 2025), setting the “model” argument to “twoway” and the “type” argument to “agreement”. For binary behavioral measures, inter-observer reliability was assessed using Cohen’s kappa coefficient. Inter-observer reliability ranged from good to excellent for continuous measures (Koo, 2015) and showed perfect agreement for binary variables (see Supplementary Tables S6 and S7 for reliability coefficients of all behavioral measures).

### Statistical analyses

To understand the effect of training approach on horse and human behavior we ran a series of (generalized) linear mixed models (GLMMs; (Baayen, 2008)) with the statistical program R (version 4.5.2) (R Core Team, 2025). Each of these models was different in their response, but had in common the training group (ABT vs. TET) included as the main predictor. In certain models we included sociability or anxiety scores as predictor as well. To be precise, the responses of affiliative and avoidance behaviors during the stroking session included sociability score as added predictor, while the responses time exploring obstacles and total refusal time during the obstacle path were analyzed with anxiety score as added predictor. Before being included in the model, these covariates were z - transformed to ease model convergence and achieve easier interpretable model coefficients (Schielzeth, 2010). As some horses had the same owner and to control for this effect, owner identity was treated as a random effect in each of these models. Three owners never completed the online questionnaire, therefore when the statistical analyses included personality scores, these data points have been excluded. Binary responses were analyzed using logistic GLMMs using the function glmer of the lme4 package (version 1.1-27.1 (Bates et al., 2015)) with family argument set to ‘binomial’; continuous responses were analyzed using linear mixed models with the lmer function of the package lmerTest (version 3.1-3; Kuznetsova et al., 2017); finally we used the glmmTMB function of the glmmTMB package (Brooks et al., 2017) to fit negative binomial models for count responses with the family argument set to ‘nbinom2’. An overview of all models can be found in the Supplementary Table S8.

Prior to fitting the linear models we inspected the responses for whether their distributions were roughly symmetrically distributed. This revealed that certain responses had a right-skewed distribution that included zeroes. To reduce skew and accommodate zeroes we applied a shifted log-transformation by adding a small value to each response before transforming it (ln(y+0.01). After fitting each model, we checked model assumptions and assessed model stability. For linear mixed models we confirmed that there were no strong deviations from assumptions of normality and homogeneity of residuals by visual inspection of qq-plots of residuals and residuals plotted against fitted values. For each model we visually confirmed that the best linear unbiased predictors (BLUPs) per level of the random effects were approximately normally distributed, which they were.

We also assessed model stability by excluding levels of random intercept effect of owner one at a time and comparing the resulting estimates with those obtained from the model based on all data (Nieuwenhuis et al., 2012), we determined that all models were of moderate to good stability.

For the linear mixed models, we tested the effect of individual fixed effects by means of the Satterthwaite approximation (Luke, 2017) using the function lmer of the package lmerTest (version 3.1-3; Kuznetsova et al., 2017) and a model fitted with restricted maximum likelihood. For the binomial and negative binomial models we tested the effect of individual predictors using the drop1 function, which can be used to compare simpler with more complex models utilizing likelihood ratio tests. We used the emmeans function of emmeans package to calculate estimated marginal means for the group effect (Lenth et al., 2023). To test for an association between groups and their crossing behavior we used a linear-by-linear association test (Agresti 2002 (p.373)) using the lbl_test function of the coin package (Hothorn 2006). We chose this approach as it takes the ordinal nature of the variables crossing behavior into account (i.e. in the order of “no pass”; “rush through”; “tense pass”; “relaxed pass”) and is thus preferred over a simpler chi-square test.

Figures were created using the ggplot2 package (Wickham 2016). Interpretation of p -values was conducted using the “language of evidence” guidelines (Muff 2022), to allow for a more nuanced way to present the results with grading from “little or no evidence” to “very strong evidence”.

### Writing

We used OpenAI, ChatGPT (February 16th 2026 version) to verify and improve readability and shorten the text. The prompt used was “Improve the readability and shorten this scientific section without changing any content and style”. The generated version was then compared to the original version, and the authors edited specific sentences in the original version when appropriate.

## Results

### Stroking session

During the stroking session, only one horse exhibited a brief sleeping episode, while 13 horses showed restlessness episodes. There was strong evidence that ABT horses were more likely to have restless episodes than TET horses (ABT: 50.1 ± 11.6%; TET: 9.90 ± 8.1%; *χ*^2^_(1)_ = 8.471, *p* < 0.01). But, we found no evidence that ABT and TET horses differed in their time spent in alert (ABT: 2.01 ± 0.83; TET: 1.60 ± 0.81; *t*_(30.98)_ = -0.363, *p* = 0.719) or resting states (ABT: 404.54 ± 16.69; TET: 419.73 ± 23.09; *t*_(33.14)_ = 0.658, *p* = 0.515).

When the brush was introduced, there was moderate evidence that horses trained with ABT were more likely to explore it compared to TET horses (ABT: 64.2 ± 11.3%; TET: 24.0 ± 11.6%; *χ*^2^_(1)_ = 6.453 *p* < 0.05). There was no evidence that the count of avoidance behaviors differed between groups (ABT: 1.65 ± 0.558; TET: 1.95 ± 0.66; *χ*^2^_(1)_ = 0.176 *p* = 0.674), nor that there was a relationship between sociability and avoidance behavior (1.79 ± 0.56; *χ*^2^_(1)_ = 1.430 *p* = 0.232). In contrast, we found moderate evidence that affiliative behaviors were more frequent in ABT horses than in TET horses (ABT: 10.36 ± 2.01; TET: 5.14 ± 1.09; *χ*^2^_(1)_ = 5.724, *p* < 0.05), and there was very strong evidence for a positive relationship between sociability and the frequency of affiliative behaviors (7.29 ± 1.06; *χ* ^2^_(1)_ = 11.842, *p* < 0.001).

### Obstacle path

There was no evidence that completion time for the obstacle path differed between groups (ABT: 4.42 ± 0.205; TET: 4.80 ± 0.207; *t*_(41)_ = 1.359, *p* = 0.182), indicating comparable task efficiency across training methods.

However, marked differences emerged in horses’ behavioral responses during task execution. We found moderate evidence that ABT horses spent more time exploring the obstacles than TET horses (ABT: 2.68 ± 0.65; TET: 0.31 ± 0.67; *t*_(32.06)_ = -2.515, *p* < 0.05, Fig. 4), while controlling for anxiety scores with no evidence that anxiety scores affect exploration behavior (Anxiety score: 1.49 ± 0.47; *t*_(30.04)_ = 0.288, *p* = 0.775).

**Figure 4:**
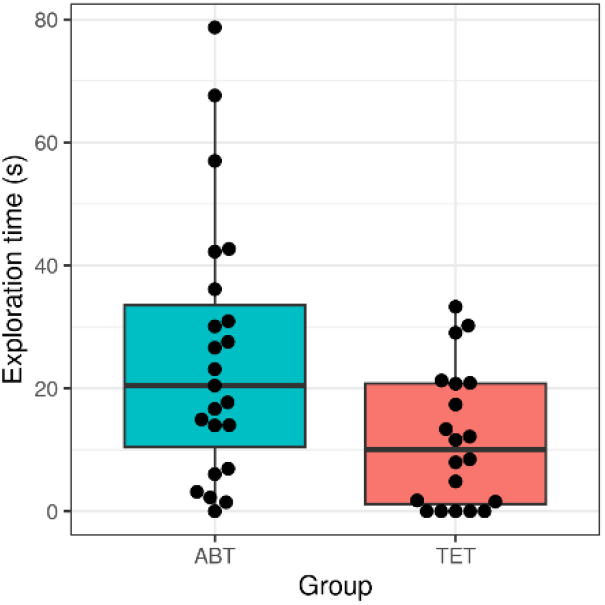
Time spent exploring obstacles during the obstacle path by horses trained with assent-based (ABT) and traditional (TET) training protocols.

Refusal time, defined as the time spent displaying behaviors that did not contribute to crossing the obstacle, was used as an indicator of compliance. We found moderate evidence that ABT horses spent less time in refusal compared to TET horses (ABT: 0.83 ± 0.76; TET: 3.37 ± 0.79; *t*_(29.59)_ = 2.336, *p* < 0.05), but we found no evidence for a relationship between refusal time and anxiety score (Anxiety score: 0.02 ± 0.54; *t*_(32.47)_ = 0.041, *p* = 0.968). Furthermore, we found strong evidence that fewer ABT horses showed proactive refusals (e.g., rearing, moving backward or sideways) than TET horses (ABT: 43.5 ± 10.3%; TET: 85.0 ± 7.9%; *χ*^2^_(1)_= 8.264, *p* < 0.01, Fig. 5).

**Figure 5:**
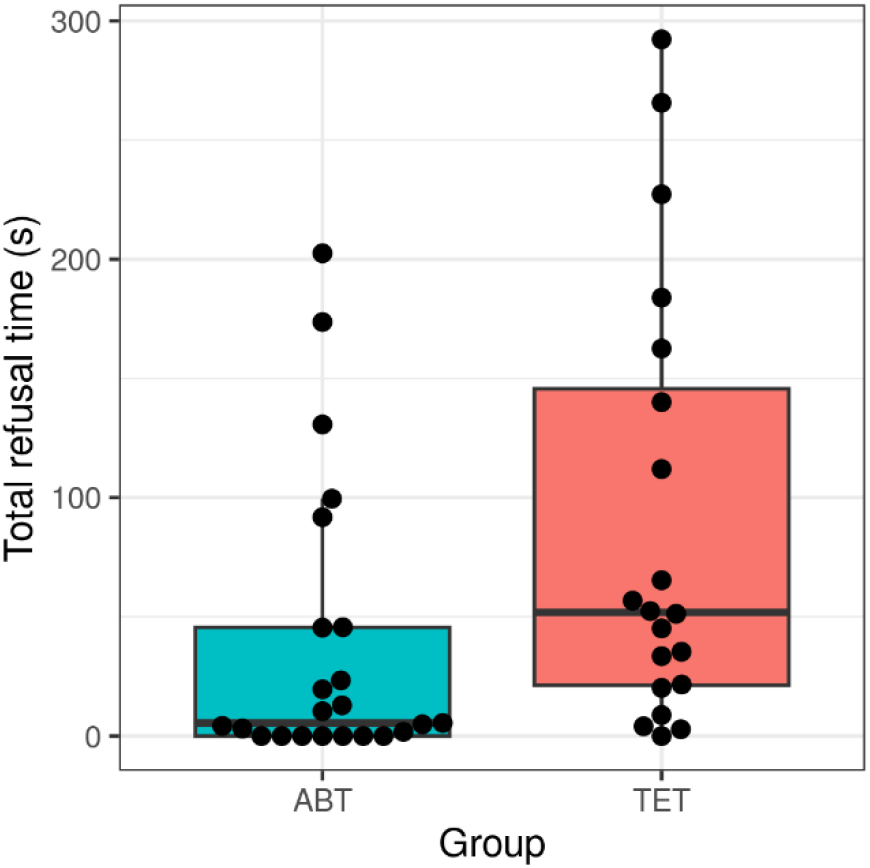
Total duration of refusal behaviors during the obstacle path in horses trained with assent-based (ABT) and traditional (TET) training protocols.

We found strong evidence for an association between groups and crossing behavior quality (*z* = 2.846, *p* < 0.01, Fig. 6). ABT horses showed a higher frequency of relaxed pass-throughs (ABT: 62, TET: 39), whereas TET horses more frequently exhibited tense crossings (ABT: 1, TET: 9), rushed movements (ABT: 5, TET: 8), or complete failure to cross (ABT: 1, TET: 4).

**Figure 6:**
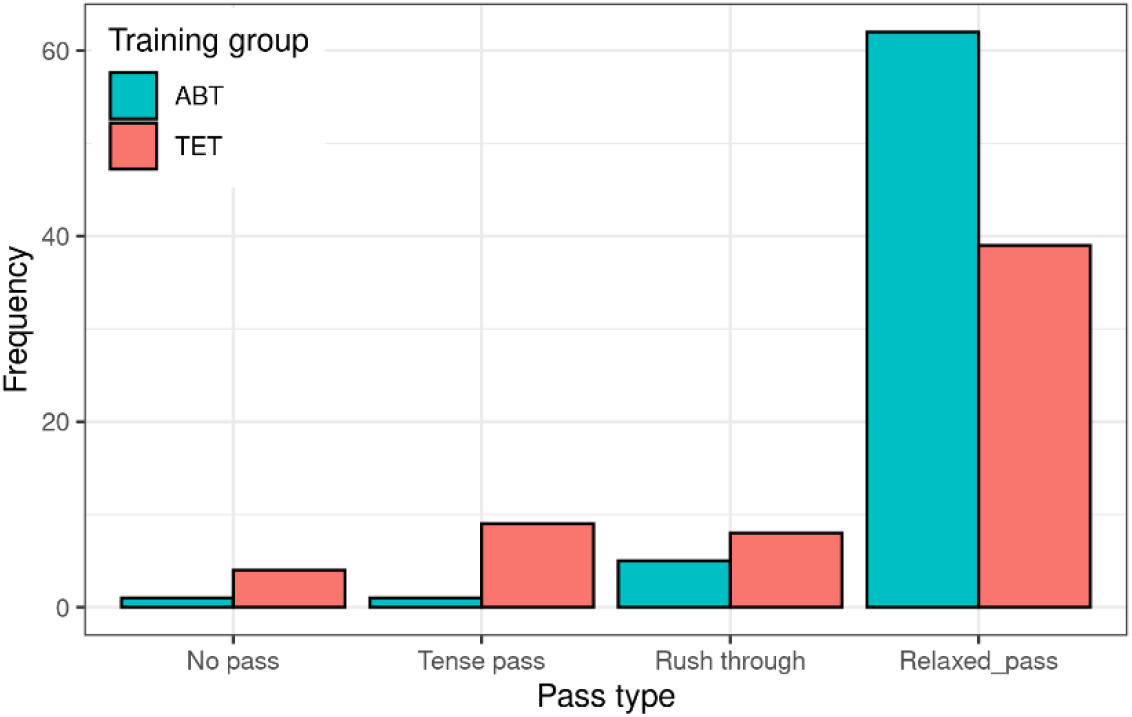
Frequency distribution of crossing behaviors in horses trained with assent-based (ABT) and traditional (TET) training protocols

Finally, we found that owner behavior also differed between groups. While almost all owners pulled on the rope occasionally, we found very strong evidence that owners in the TET group pulled on the rope for longer (ABT: 0.97 ± 0.53; TET: 3.62 ± 0.53; *t*_(41)_ = 3.677, *p* > 0.001). We found moderate evidence that TET owners used vocal praise less frequently (ABT: 3.63 ± 1.26; TET: 1.38 ± 0.53; *χ*^2^_(1)_= 4.541, *p* < 0.05), and although physical praise occurred at low levels in both groups, we found weak evidence that TET owners used slightly more physical praise compared to owners in the ABT group (ABT: 0.36 ± 0.13; TET: 0.67 ± 0.13; *χ*^2^_(1)_= 2.935, *p* = 0.08).

**Table 1.** Summary of behavioral outcomes associated with assent-based (ABT) and traditional equine training techniques (TET) across experimental contexts. “=” indicates no detectable difference.

| Experimental context | Behavioral outcome | Results |
| --- | --- | --- |
| Stroking session | Attitude | Restlessness: ABT > TET |
|  |  | Alert: ABT = TET |
|  |  | Rest standing: ABT = TET |
| Stroking session | Brush exploration | ABT > TET |
| Stroking session | Affiliative behaviors | ABT > TET |

| ***Experimental context*** | ***Behavioral outcome*** | ***Results*** |
| --- | --- | --- |
| *Stroking session* | *Avoidance behaviors* | *ABT = TET* |
| *Obstacle path* | *Completion time* | *ABT = TET* |
| *Obstacle path* | *Overall refusal time* | *ABT < TET* |
| *Obstacle path* | *Proactive refusal occurrences* | *ABT < TET* |
| *Obstacle path* | *Obstacles exploration* | *ABT > TET* |
| *Obstacle path* | *Crossing behavior quality* | *No pass: ABT < TET* |
|  |  | *Rush through: ABT < TET* |
|  |  | *Tense pass through: ABT < TET* |
|  |  | *Relaxed pass through: ABT > TET* |
| *Obstacle path* | *Human behavior* | *Pressure applied to the rope: ABT < TET* |
|  |  | *Vocal praising: ABT > TET* |
|  |  | *Physical praising: ABT = TET* |

## Discussion

This study offers novel insights into how assent-based equine training techniques (ABT) may shape the human – horse relationship compared to more traditional training techniques (TET). Overall, the findings suggest that while performance did not differ across training styles in the obstacle path, behavioral responses, emotional expression, and interaction quality varied significantly in favor of the assent-based training group across both tests.

The stroking session imposed a standardized, low-demand interaction scenario. Despite the passive nature of the context, ABT horses remained behaviorally engaged i.e. were more restless and displayed significantly more positive human-directed behaviors, such as grooming and sniffing. Moreover, when the brush was introduced, ABT horses actively explored and interacted with it, suggesting that they were not passive recipients of human touch, but rather participated in shaping the interaction. This contrasts with the typical interpretation of grooming sessions as one-directional interactions in which the human acts upon the horse rather than a reciprocal exchange (Lansade, 2019). These results could suggest that ABT horses develop a stronger horse–owner attachment bond (Lundberg, 2020) or/and that ABT horses are more confident to express behaviours.

Interestingly, sociability scores were positively associated with the expression of affiliative behaviors, but not with avoidance behaviors. This finding suggests that owner-perceived sociability reflects a horse’s general propensity to engage positively in social interactions rather than a broader tendency to express social behaviors of all types. Owner-perceived sociability was evaluated through Likert-scale ratings of behaviors such as initiating or joining play when solicited, being sought out as a companion by others, and seeking the companionship of conspecifics. Accordingly, horses rated as more sociable by their owners displayed more affiliative behaviors during the stroking session, regardless of training group, supporting the interpretation of sociability as a stable personality trait associated with social motivation and positive engagement with others (Brubaker, 2021). Sociability scores did not differ between the two training groups, indicating that baseline levels of social motivation were comparable. Nevertheless, ABT horses expressed significantly more affiliative behaviors than TET horses. This suggests that while sociability may provide the underlying predisposition for positive social interaction, the quality of the human–horse relationship influences how readily this predisposition is expressed in interactions with the owner. In other words, sociability appears to set a baseline for affiliative tendencies, whereas relationship experiences may modulate the extent to which these behaviors are displayed in a given context. The absence of a relationship between sociability and avoidance behaviors further supports this interpretation. Avoidant responses may be influenced less by stable social disposition and more by contextual factors, such as the horse’s perception of the interaction, previous experiences with the handler, or momentary emotional state. Consequently, the increased expression of affiliative behaviors observed in ABT horses cannot be attributed solely to differences in personality, but instead suggests that the training approach may have contributed to the development of a more positive human–horse relationship.

Interestingly, ABT horses showed a higher occurrence of restless episodes during the stroking session, than TET horses, which could be in line with showing more communicative behaviors. Alternatively, as most ABT owners expressed that their horses were not accustomed to being tied, as required by the standardized protocol, this could also be interpreted as increased arousal in a new situation. Nevertheless, the fact that the ABT horses showed more affiliative behaviours during the test situations, suggests that they were not completely uncomfortable. Future studies might consider a refinement of the stroking session to give complete freedom of movement and an active role to the horses: a scenario where the horse is free to approach the owner on its own terms might be more reflective of the HHR horse’s perception.

To complement these findings and provide a more comprehensive assessment of the HHR, the obstacle path was designed to evaluate horse behavior in a challenging context involving novel stimuli and reliance on the owner. The horses’ anxiety scores did not significantly predict the behavioral outcomes, supporting the interpretation that the training method itself, rather than individual emotional predisposition, played a central role in shaping the horse’s behavioral responses during the novel and potentially challenging tasks.

Although both groups completed the obstacle path in comparable times, their behavioral responses were markedly different. ABT horses dedicated more time to exploratory behavior and less to refusal behaviors, particularly proactive refusal, suggesting that the presence of their human partner may serve as a secure base that encourages investigation and reduces avoidance as has been observed across many other species including dogs (Horn, 2013)), wolves (Lenkei, 2020) and cats (Takeda, 2024). This interpretation is in line with the results from the stroking session and resonates with attachment theory (Mikulincer, 2003): the presence of an attachment figure can provide a sense of safety and reduce stress, thereby facilitating exploration and interaction with the environment (Bowlby, 1979). Moreover, ABT horses were also significantly less likely to show proactive refusal in the obstacle task than TET horses. However, while the horses in the ABT group seemed more interested and relaxed, it is difficult to disentangle if that is due to the experienced training style, which may or may not have helped to forge a good relationship with the owner, or due to differences in owner’s behavior and personality. Also the active refusals, which can be potentially dangerous for the human handler, might be triggered by insecurity of the owner.

In regard to the owners’ behavior we found that almost all owners (in both groups) pulled on the rope occasionally, but while TET owners relied on pulling for significantly longer durations, owners in the ABT group tended to praise verbally small steps more frequently. These differences in the owners interaction style could either reflect the different training philosophies or indicate that people with similar personalities use similar training styles.

In line with this, leading a horse through an obstacle path may trigger different emotional states in humans based on their personalities. Horses have a well-known ability to perceive human emotions and intentions as demonstrated by the famous Clever Hans, who was able to give correct answers to mathematical questions by reading the microscopic signals in the face of the questioning person (Samhita,2013; Pfungst, 1911). Moreover, in another study, it was shown that horses’ heart rate increased when a potentially dangerous event about to happen was communicated to their owners (Keeling at al. 2009).

In general, it is difficult to determine whether differences in horses’ behaviour reflect training style, the quality of the caregiver–horse relationship, or owner personality. These factors are likely interdependent: training style shapes the relationship, while owner personality influences both the methods adopted and the degree of security conveyed to the animal. Nevertheless, other studies have detected no differences in stress responses between the handling by the owner and a stranger (Ijichi, 2018) nor found that only the owner can evoke a “safe haven” effect in horses (Lundberg, 2020) suggesting that owner personality cannot fully explain the horses’ behaviour. Moreover, studies in zoos and wild animals’ management have demonstrated that providing animals with choice opportunities enhances also safety for the caregiver (Brando, 2023), which was seen in our study in the lower count of active refusals in the ABT compared to TET horses. Finally, assent-based methods are practiced only by a minority of horse’ owners and one might argue that adopters likely share certain attitudes and personality profiles that then influence the owners’ behaviors and in turn the horses’ behavior. However, the TET owners were not professional riders and appeared welfare-oriented since most housed their horses in paddocks with conspecifics, suggesting that differences in attitudes and personalities might not have been that different between our two groups.

Overall, the results lend support to our hypothesis that assent-based training techniques promote a more exploratory, confident, and communicative interaction style in horses, which may in turn strengthen the quality of the HHR. However, due to the cross-sectional design, causal inferences cannot be drawn, and past experiences, type of owner-horse interactions and owner personality and competence may have contributed to the observed differences. Furthermore, whether the observed effects are specifically related to the assent-based protocols or rather to the use of positive reinforcement could not be distinguished in this study. Positive reinforcement in itself has been found to have positive effects on horse–human interactions: it increases horse interest in humans (Sankey, 2010), motivation to participate in training sessions and exhibition of exploratory behavior (Innes, 2008).

Despite these limitations, the present findings contribute to a growing body of evidence highlighting the relevance of training techniques in shaping the HHR and in promoting equine welfare. From a welfare perspective, the findings provide preliminary empirical support for the idea that assent-based training approaches may enhance horses’ perceived control and willingness to engage with humans. The combination of positive human-directed behaviors in the stroking session, reduced refusal and more frequent relaxed crossing behaviors in the obstacle path and increased exploration in both tasks, suggest that interactions may be experienced as less threatening and more predictable for ABT horses. Importantly, the absence of differences in task completion time in the obstacle path addresses concerns that granting horses greater control might compromise effectiveness. In this study, assent-based training was associated with different behavioral strategies rather than reduced performance and compliance.

## Supporting information

ObstaclePath_data

StrokingSession_data

## Data availability

The datasets generated and analysed during the current study, containing the behavioral measurements used for the statistical analyses are provided as .xlsx files in the Supplementary Information. All other data supporting the findings of this study are included in this published article and its supplementary information files. Additional raw material, including the original BORIS coding files, is available from the corresponding author on reasonable request.

## Author contribution

ET was involved in design of the methods, data collection, data analysis, and writing of the manuscript. FR was involved in design of the methods, writing of the manuscript and supervision. RF was involved in data analysis. All authors reviewed the manuscript.

## Acknowledgements

We would like to thank the Domestication Lab members for all the support and feedback; the horse-owners who participated with their horses; the ranch-owners who gave full time and environment availability and participated in the planning of the tests.

