## Supplementary Information for "Beyond compliance: evaluating the impact of assent-based training on equine behavior and the human-horse relationship"

S1: list of horse participants and their general information

| Name | Sex | Age | Breed | Housing | Social life | Training style |
|---|---|---|---|---|---|---|
| Asmar | Gelding | 13 | Arabian | Paddock 24h | Permanent herd (n =4) | ABT |
| Bali | Female | 7 | Haflinger | Paddock 24h | / | ABT |
| Ester | Female | 14 | Mix | Paddock 24h | Permanent herd (n =12) | ABT |
| Garibaldi | Gelding | 10 | Mix | Paddock 24h | Permanent couple | ABT |
| Harry | Gelding | 15 | Arabian | Paddock 24h | Herd (n =35) | ABT |
| Iris | Female | 19 | Quarter Horse x Haflinger | Paddock 24h | Permanent couple | ABT |
| Kim | Female | 9 | Irish mix | Paddock 24h | Permanent herd (n =4) | ABT |
| Maja | Female | 11 | Noriker | Paddock 24h | Permanent herd (n =3) | ABT |
| Makha | Female | 7 | Criollo x Paint Horse | Paddock | Permanent couple | ABT |
| Myricae | Female | 9 | Merence | Paddock 24h | Permanent herd (n =3) | ABT |
| Ofelia | Female | 25 | Sella Italiana | Paddock 24h | Permanent couple | ABT |
| Ori | Gelding | 7 | Paint Horse | Paddock 24h | Herd (n =35) | ABT |
| Pandela | Female | 14 | Anglo-arabian | Paddock 24h | Permanent couple | ABT |
| Pemba | Female | 13 | Akhal Teke | Paddock 24h | Herd (n =7/10) | ABT |
| Saleg | Gelding | 8 | Belgian Throughbred | Paddock 24h | Herd (n =35) | ABT |
| Fe | Gelding | 16 | Argentine | Paddock 24h | Permanent herd (n =4) | ABT |
| Seventh | Gelding | 9 | Irish cob | Paddock 24h | Permanent herd (n =3) | ABT |
| Shasir | Gelding | 15 | Mix | Paddock 24h | Herd (n =4) | ABT |
| Sofi | Female | 24 | Quarab | Paddock 24h | Herd (n =35) | ABT |
| Tequila | Female | 18 | Mix | Paddock 24h | Herd (n =35) | ABT |
| Thiago | Gelding | 14 | Arabian Mix | Paddock 24h | Permanent herd (n =12) | ABT |
| Utopia | Female | 11 | Mix | Paddock 24h | Permanent herd (n =12) | ABT |
| Silver | Gelding | 9 | Mix | Paddock 24h | Permanent herd (n =4) | ABT |
| Spirit | Gelding | 13 | Mix | Paddock 24h | Permanent herd (n =12) | ABT |
| Africa | Female | 10 | Paint Horse | Box + paddock | / | TET |
| Calliope | Female | 12 | Sella italiana | Box + paddock | / | TET |
| Casper | Gelding | 10 | Spanish x Appaloosa | Paddock + box at night | / | TET |
| Cervinia | Female | 13 | Bardigiano | Paddock + box at night | / | TET |

| | | | | | | |
|---|---|---|---|---|---|---|
| Chilly | Female | 6 | Quarter Horse | Box + paddock | / | TET |
| Doc | Gelding | 16 | Appaloosa | Paddock + box at night | Permanent couple | TET |
| Jill | Gelding | 13 | Paint Horse x Quarter Horse | Paddock + box at night | Permanent couple | TET |
| Linda | Female | 10 | Quarter Horse | Paddock 24h | / | TET |
| Miss | Female | 9 | Paint Horse | Paddock 24h | / | TET |
| No joke | Gelding | 9 | Dutch | Box + paddock | / | TET |
| Parny | Female | 8 | English Throughbred | Paddock + box at night | Permanent couple | TET |
| Persefone | Female | 6 | Connemara | Paddock + box at night | / | TET |
| Pico | Gelding | / | / | Paddock 24h | / | TET |
| Red | Gelding | 16 | Quarter Horse Mix | Paddock 24h | Permanent couple | TET |
| Rope | Female | 8 | Quarter Horse | Box | / | TET |
| Sajib | Stallion | 30 | Arabian Mix | Paddock | / | TET |
| Shakira | Female | 5 | Quarter Horse | Box + paddock | / | TET |
| Spartaco | Gelding | 17 | Spanish Mix | Paddock 24h | Herd (n =2/3) | TET |
| Speedy | Gelding | 17 | Appaloosa | Paddock 24h | / | TET |
| Topper | Gelding | 13 | Quarter Horse Mix | Paddock 24h | Permanent couple | TET |
| Winky | Gelding | 12 | Paint Horse | Paddock 24h | Permanent couple | TET |

S2: survey items extracted from the HPQ of Lloyd at al. (2008) and included in the personality section of the questionnaire.

| Item ID | Question | Personality trait |
|---|---|---|
| 1 | Try to readily avoid others or outside disturbances | ANXIOUSNESS |
| 2 | Hesitates to act alone, seeks reassurance from others | ANXIOUSNESS |
| 3 | Initiates play and joins in when play is solicited | SOCIABILITY |
| 4 | Sought out as a companion by others | SOCIABILITY |
| 5 | Seeks companionship of others | SOCIABILITY |
| 6 | Shows restraint in posture and movement; carries the body stiffly, which suggests a shrinking tendency, as if to pull back and be less conspicuous | ANXIOUSNESS |
| 7 | Does not trust others readily (humans and horses), trusts few individuals | ANXIOUSNESS |

For each item, respondents indicated their likelihood on a five-point Likert scale ranging from very unlikely to very likely. Responses were coded using a symmetric scoring scheme (single score, SC), where very unlikely = −2, unlikely = −1, neutral = 0, likely = +1, and very likely = +2.

Personality trait scores were computed by summing the SC values across all items associated with the respective trait (e.g., Anxiousness = SC(1) + SC(2) + SC(6) + SC(7); Sociability = SC(3) + SC(4) + SC(5)), with higher scores indicating stronger expression of that trait personality section of the survey.

S3: training section of the survey.

| Item ID | Question | Scale | Training concept |
|---|---|---|---|
| 1.1 | I use pressure and release when I want to teach my horse something new | Never - Always | R+ |
| 1.2 | When I want my horse to move forward I use my driving aids (leg pressure and/or rein pressure) | Never - Always | R- |
| 1.3 | When my horse performs correctly an exercise, I give him a break | Never - Always | Buffer |
| 1.4 | If my horse doesn't respond to my aids I emphasize them by intensifying the pressure | Never - Always | R- |
| 1.5 | I sometimes gently touch my horse with the whip to clarify what I want it to do | Never - Always | R- |
| 1.6 | I allow my horse to eat grass after the exercise sessions | Never - Always | Buffer |
| 1.7 | If my horse performs a behavior I didn’t ask for, I correct it | Never - Always | R- |
| 1.8 | I give my horse a treat when he performs a correct behavior | Never - Always | R+ |
| 1.9 | When I want to teach my horse something new I wait until the he/she offers the behavior by itself | Never - Always | R+ |
| 1.10 | I play with my horse in order to lure out new behaviours for which I can give rewards | Never - Always | R+ |
| 1.11 | If my horse refuses to move forward I lure with a treat | Never - Always | R+ |

| | | | |
|---|---|---|---|
| 1.12 | If my horse performs a behavior I didn't ask for, I ignore it | Never - Always | R+ |
| 1.13 | I do exercises to help my horse become more athletic | Never - Always | Buffer |
| 1.14 | I use my riding aids (reins pressure and/or leg pressure,...) to help my horse keep the proper posture | Never - Always | TET |
| 1.15 | I invite my horse to chase objects like balls to help him develop agility | Never - Always | ABT |
| 1.16 | I do exercises with my horse to help him regulate its energies | Never - Always | Buffer |
| 2.1 | If my horse is not comfortable with a procedure, I use a consent protocol (I teach him signals that he/she will use to tell me yes or no) | Never - Always | ABT |
| 2.2 | If my horse doesn't respect my personal space (for example tends to push me), there is the risk he/she is trying to establish dominance over me. | Strongly disagree – strongly agree | TET |
| 2.3 | If I see an adult (not green) horse refusing to offer its hooves, I consider the possibility that he might have balance issues | Strongly disagree – strongly agree | ABT |
| 2.4 | If my horse struggles with separation anxiety (from his pony friend), it is a good idea to bring also his friend to the arena during training sessions. | Strongly disagree – strongly agree | ABT |
| 2.5 | If my horse tries to avoid saddling by side stepping, it's my priority to correct it to prevent the chance it becomes a habit | Strongly disagree – strongly agree | TET |
| 2.6 | If I see an adult (not green) horse refusing to offer his hooves, I consider the possibility that he is challenging the handler | Strongly disagree – strongly agree | TET |
| 2.7 | If I sometimes leave some choices to my horse (for example do not ride him if he moves away from the mounting block), he might become difficult to handle | Strongly disagree – strongly agree | TET |

Items assessed different training concepts (positive reinforcement [R+], negative reinforcement [R−], assent-based training [ABT], and traditional training [TET]) using five-point Likert scales (e.g., never to always, strongly disagree to strongly agree). The items related to R+ and R- were extracted from the study of Lundberg at al. (2020).

Responses were converted to item-level scores (SC) on a scale from 0 to ±4. Items reflecting R+ and ABT were scored positively (0 to +4), whereas items reflecting R− and TET were scored negatively (0 to −4). A composite training score (CTS) was calculated for each participant as the sum of all item-level scores, with higher values indicating greater alignment with R+/ABT practices and lower values indicating greater alignment with R−/TET practices.

Participants had been previously assigned, based on a preliminary phone interview, to ABT or TET training group. Mean training scores were then calculated separately for each group. The midpoint between the two group means was used as a cut-off value to evaluate whether participants' questionnaire-based scores were consistent with their initial group classification. All participants CTS confirmed the training group assigned through the phone interview.

S4: ethogram for the stroking session

| Category | Behavior | Definition |
|---|---|---|
| **ATTITUDE** | Stand Alert | Rigid stance with the neck elevated and the head oriented toward the object or animal of focus. The ears are held stiffly upright and forward, and the nostrils might be slightly dilated. (McDonnell 2003) |
| | Rest Standing | Standing inactive in a relaxed posture, usually with head slightly lowered, eyes partly or nearly closed and often bearing weight on three legs (one hind leg slightly flexed). Lips relaxed and ears rotate laterally. (McDonnell 2003) |
| | Stand Sleep | Eyes closed, head lowered below the back, weight all on three legs (one hind leg stand on the top of the hoof), ears rotate laterally. (McDonnell 2003) |
| | Restless | The horse is constantly in movement, either walking in circle around the owner or trying to walk away. |
| | Exploration | Nose or mouth touching the brush, lips or tongue manipulating the brush, biting, chewing, licking, nibbling and sniffing. |
| **AVERSIVE BEHAVIORS** | Threat | Lunge or rapid movement of the head toward the owner with both ears backwards. |
| | Kicks | Rapid rise of the hind leg in direction of the owner to hurt him. |
| | Rear | Stand up on just back legs, may paw or strike forward with the front legs. |
| | Scream | Forceful expulsion of air through the nostrils incidentally preceeded by a raspy inhalation sound. |
| | Snort | The horse is producing a high pitch, alarming sound |
| | Avoid | Movement of the head or body in the opposite direction of the owner without moving the legs away. |

| | | |
|---|---|---|
| | Step aside<br>Stomp | Movement of one or more of the limbs on the opposite direction of the experimenter<br>Vivid step on the floor with one leg without movement. |
| **HUMAN-DIRECTED BEHAVIORS** | Groom<br>Scratch<br>Look at<br>Smell | Gently nibbling, biting, licking or rubbing on the owner.<br>Movement of gently rubbing of the head of the body against the owner.<br>Head and gaze directed toward the owner.<br>Approach of the nose close by the owner. |

S5: ethogram for the obstacle path

| Category | Behavior | Definition |
|---|---|---|
| **CROSSING TIME** | Crossing time | Starting when the horse overcomes the starting pole with the forelegs and ending when the horse overcomes the end pole with the hindlegs. |
| **EXPLORATION** | Exploration | Nose, mouth, forelegs or other body parts actively touching or moving the object, lips or tongue manipulating the object, biting, chewing, licking, nibbling and sniffing. |
| **REFUSAL BEHAVIORS** | Stationary refusal<br>Proactive refusal | Any behavior not contributing to crossing or exploring the object, while standing. The horse is still but not contributing to cross the object. (Ijichi, 2008)<br>Any refusal behaviour that involved movement, thus excluding stationary refusal: moving backwards, sideways, forwards but away from the object. (Ijichi, 2008) |
| **OWNER BEHAVIOR** | Vocal praising<br>Pressure applied to the rope<br>Physical praising | The owner is emitting any sound intended to praise the horse. It includes praising phrases like “Good boy!” or “Well done!”. Encouraging cues like “come” or “let’s go!” are not included.<br>The owner is pulling the rope, which is tense.<br>The owner pet, scratch or cuddle the horse. |
| **CROSSING BEHAVIOR QUALITY** | Rush through | The horse passes through/on the obstacle with a run (trot or gallop stride). He might pull the rope while rushing. |

| | | |
|---|---|---|
| | Tensed pass through | The horse increases the gait while overcoming the obstacle, with neck elevated, ears held stiffly, dilated nostrils and tensed muscles. He might pull the rope while passing. |
| | Relaxed pass through | The horse overcomes the obstacle without increasing the gait, in a relaxed posture with head lowered. The horse does not pull the rope while passing through. |
| | No pass through | The horse doesn't overcome the obstacle within the 5 minutes. |

S6: Inter-observer reliability coefficients for continuous (intraclass correlation coefficients, ICC) behavioral measures of the Stroking Session (SS) and Obstacle Path (OP).

| Behavioral test | Behavioral measure | Coefficient | p-value | 95% CI |
|---|---|---|---|---|
| SS | Active states (alert, rest, sleeping, restless) | **.985** | <0.000001 | .971 - .992 |
| SS | Human directed behaviors (avoidance and affiliative) | **.878** | <0.000001 | .722 - .950 |
| OP | Exploration | **.996** | <0.000001 | .989 - .999 |
| OP | Crossing time | **1** | <0.000001 | 1 - 1 |
| OP | Total refusal time | **.984** | 0.00000131 | .923 - .997 |
| OP | Owner pressure on the rope | **.931** | 0.000115 | .720 - .985 |
| OP | Crossing behavior quality | **.839** | <0.000001 | .664 - .927 |
| OP | Vocal praising | **.896** | 0.000785 | .565 - .978 |

S7: Inter-observer reliability coefficients for binary (Cohen's kappa, κ) behavioral measures of the Stroking Session (ss) and Obstacle Path (OP).

| Behavioral test | Behavioral measure | Coefficient | p-value |
|---|---|---|---|
| SS | Brush exploration | **1** | 0.00157 |
| OP | Proactive refusals | **1** | 0.00468 |
| OP | Physical praisings | **1** | 0.000785 |

S8: statistical models

| Behavioral test | Behavioral measure | Response type | Predictor(s) | Approach |
|---|---|---|---|---|
| SS | Restlessnes episodes | binomial | Group | Mixed logistic regression |
| SS | Time standing alert | continuous (shifted log-transformation) | Group | Linear mixed model |
| SS | Time resting | continuous | Group | Linear mixed model |
| SS | Brush exploration | binomial | Group | Mixed logistic regression |

| | | | | |
|---|---|---|---|---|
| SS | Avoidance behavior | count | Group + Sociability score | Negative binomial mixed model |
| SS | Human directed behavior | count | Group + Sociability score | Negative binomial mixed model |
| OP | Crossing time | continuous (log-transformation) | Group | Linear mixed model |
| OP | Time exploring obstacles | continuous (shifted log-transformation) | Group + Anxiety score | Linear mixed model |
| OP | Total refusal time | continuous (shifted log-transformation) | Group + Anxiety score | Linear mixed model |
| OP | Refusal behavior | binomial | Group | Mixed logistic regression |
| OP | Crossing behavior quality | count | Group | Linear-by-linear association test |
| OP | Owner pressure on rope | continuous (shifted log-transformation) | Group | Linear mixed model |
| OP | Vocal praise | count | Group | Negative binomial mixed model |
| OP | Physical praise | binomial | Group | Mixed logistic regression |
